# Tracking Subclonal Mutation Frequencies Throughout Lymphomagenesis Identifies Cancer Drivers in Mouse Models of Lymphoma

**DOI:** 10.1101/157800

**Authors:** Philip Webster, Joanna C. Dawes, Hamlata Dewchand, Katalin Takacs, Barbara Iadarola, Bruce J. Bolt, Juan J. Caceres, Jakub Kaczor, Laurence Game, Thomas Adejumo, James Elliott, Kikkeri Naresh, Ge Tan, Gopuraja Dharmalingam, Alberto Paccanaro, Anthony G. Uren

**Author notes:** Denotes shared first authorship (Philip Webster, Joanna C. Dawes).

## Abstract

Determining whether recurrent but rare cancer mutations are bona fide driver mutations remains a bottleneck in cancer research. Here we present the most comprehensive analysis of retrovirus driven lymphomagenesis produced to date, sequencing 700,000 mutations from >500 malignancies collected at time points throughout tumor development. This enabled identification of positively selected events, and the first demonstration of negative selection of mutations that may be deleterious to tumor development indicating novel avenues for therapy. Customized sequencing and bioinformatics methodologies were developed to quantify subclonal mutations in both premalignant and malignant tissue, greatly expanding the statistical power for identifying driver mutations and yielding a high-resolution, genome wide map of the selective forces surrounding cancer gene loci. Screening two BCL2 transgenic models confirms known drivers of human B-cell non-Hodgkin lymphoma, and implicates novel candidates including modifiers of immunosurveillance such as co-stimulatory molecules and MHC loci. Correlating mutations with genotypic and phenotypic features also gives robust identification of known cancer genes independently of local variance in mutation density. An online resource http://mulv.lms.mrc.ac.uk allows customized queries of the entire dataset.

## INTRODUCTION

The majority of verified human cancer genes have been identified by translocation breakpoints, recurrent exonic mutations, and focal copy number aberrations (Futreal et al. 2004). Increasing cohort sizes of human tumor sequencing has revealed large numbers of rare clonal mutations, however their contribution to disease is more difficult to prove due to a lack of statistical power, giving rise to false positives and negatives (Lawrence et al. 2013). It is similarly challenging to determine how deregulation of intact open reading frames by non-coding mutations, large-scale copy number alterations and epigenetic mechanisms contributes to disease. The data available to identify cancer drivers from tumor sequencing studies could be increased through the study of subclonal mutations in both premalignant samples as well as mature tumors, however this requires quantification of both in numbers sufficient to demonstrate that selection has taken place.

Murine leukemia virus (MuLV) induced lymphoma is an ideal model system to study the extent to which subclonal mutations can be used to identify cancer drivers. Infection of newborn mice with MuLV causes a systemic lifelong viremia whereby viral integrations deregulate and truncate nearby genes by diverse mechanisms. Fusion transcripts, virus LTR enhancer elements acting on endogenous promoters and replacement of 3’ UTR sequences can increase expression, whilst disruption of endogenous transcripts, transcriptional interference or methylation of DNA flanking integrations can decrease expression (Berns 2011). These mutations eventually give rise to hematological malignancies, and a high proportion of the recurrently mutated loci correspond to known drivers of human lymphoid malignancies, as well as regulators of hematopoietic development and lymphocyte survival (Berns 2011; Ranzani et al. 2013). Historically, these screens focused on mutations present in clonal outgrowths as evidence of their role in malignancy, however recent pyrosequencing of MuLV lymphomas has also shown selection taking place within subclonal populations of cells (Huser et al. 2014). Cloning integration mutations by ligation mediated PCR requires a fraction of the sequencing coverage needed to identify other mutation types, allowing large numbers of subclonal integration mutations to be identified with unparalleled sensitivity. Furthermore, gamma retroviruses are not subject to remobilization, can integrate in any sequence context, and localized bias of the orientation of integrations can be used as a measure of selection that is independent of regional variation in integration density (Huser et al. 2014).

The fraction of rare mutations that drive cancer is largely unknowable. In this study, we use somatic insertional mutagenesis in mice as a model to demonstrate that subclonal mutations that are only rarely found as clonal mutations in advanced-stage disease can be effectively employed to identify known cancer drivers and differentiate rare disease causing mutations from passenger mutations. Using a novel insertion site cloning protocol, able to detect subclonal retroviral integrations with unprecedented sensitivity, we cloned more than 3000 clonal and 700,000 subclonal mutations across a spectrum of > 500 MuLV induced T-cell and B-cell lymphoid malignancies from two *BCL2* transgenic models over a time course of lymphomagenesis. From these we find both positive and negative selection of subclonal events throughout all stages of lymphomagenesis and that in late-stage disease both clonal and subclonal populations identify more than 100 known cancer drivers and regions implicated in non-Hodgkin lymphoma (NHL) by coding mutations, copy number aberrations and genome wide association studies (GWAS). This resource can be used to prioritize rare but recurrent mutations from human tumors for further study.

## RESULTS

### A time course of MuLV infection quantifies the transition from premalignancy to malignant lymphoma

To observe how subclonal mutations undergo selection during lymphomagenesis we generated a diverse set of B cell and T cell derived lymphoid malignancies, sacrificing animals with advanced-stage disease, as well as at a series of premalignant time points. Moloney MuLV typically results in a T-cell leukemia/lymphoma however subtype and mutation profile can be skewed by genetic background and predisposing germline alleles (Uren et al. 2008; Kool et al. 2010). To produce malignancies where mutation profiles are correlated with different phenotypes and predisposing genotypes, we generated lymphomas on two genetic backgrounds using both wild type animals and two *BCL2* transgenic models. The t(14;18)(q32;q21) *IGH/BCL2* translocation drives enforced expression of the antiapoptotic protein BCL2 and is one of the earliest and most common initiating mutations of follicular lymphoma (FL) and diffuse large B-cell lymphoma (DLBCL). Overexpression of BCL2 is also frequently observed in B-cell chronic lymphocytic leukemia (CLL).

Newborns were infected with MuLV by intraperitoneal injection. A cohort of Vav-BCL2 transgenic animals (expressing high levels of human *BCL2* from a Vav promoter (Ogilvy et al. 1999)) and wild type littermates were generated on a C57BL/6 x BALB/c F_1_ background. Emu-BCL2-22 transgenic cohorts (expressing moderate levels of human *BCL2* from an Emu enhancer (Strasser et al. 1991)) and wild type littermates were generated on both C57BL/6 x BALB/c F_1_ and C57BL/6 backgrounds. All mice developed lymphoid malignancies with latency ranging 42-300 days (Fig. 1a-c), with enlarged spleens, thymuses and lymph nodes observed in all cohorts. Disease onset was significantly accelerated by the Vav-BCL2 transgene on the F_1_ background (p=0.0001) (Fig. 1a) and by the Emu-BCL2-22 transgene on a C57BL/6 background (p=0.0163) (Fig. 1c) compared with their littermate controls. The F_1_ background developed lymphoma more rapidly than equivalent C57BL/6 cohorts (supplemental Fig. S1).

**Figure 1.**
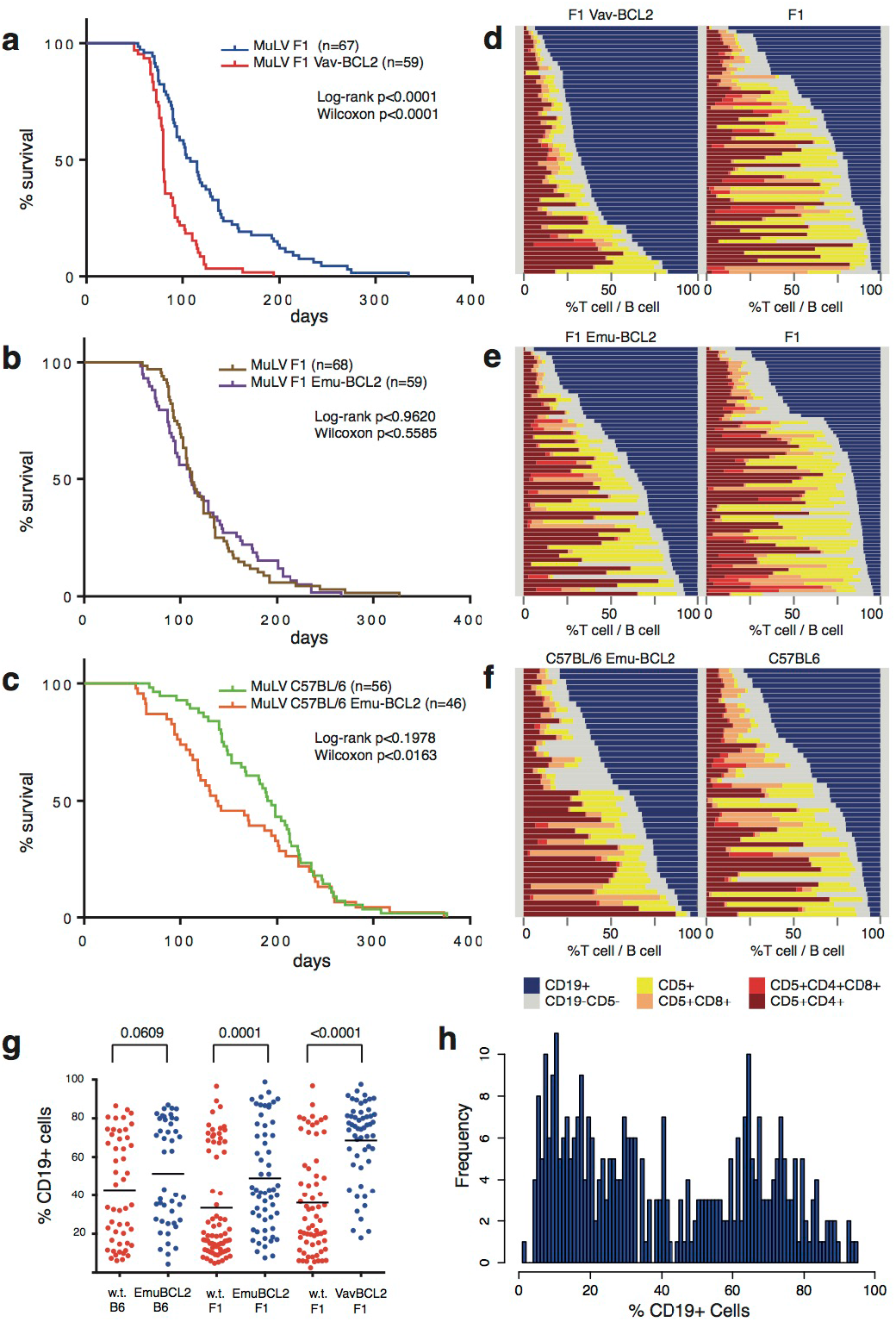
Variable latency and immunophenotype of MuLV lymphoma from wild type and BCL2 transgenic mice. a) The Vav-BCL2 transgene significantly reduced latency on an F_1_ background. b & c) The Emu-BCL2-22 transgene significantly reduces latency on a C57BL/6 background but not F_1_ background. The Emu-Bcl2-22 C57BL/6 cohort had a significantly shorter latency than wild type C57BL/6 controls and both C57BL/6 cohorts had longer latency compared with F1 equivalents. (Supplemental Fig. 1). d-f) Stacked bar charts on the right represent the immunophenotyping of spleen suspensions from each cohort. Each row represents one spleen. Colors in each row represent the proportion of B cells (blue CD19+) and T cells (yellow CD5+ CD4-CD8-, light orange CD5+ CD8+, dark orange CD5+ CD4+ CD8+ and red CD5+ CD4+) in each sample. BCL2 transgenes increase the proportion of B cells in all cohorts and the mixture of T cell lymphoma subtypes is highly variable. g) The proportion of CD19+ B cells is increased by both BCL2 transgenes (h) Histogram of all CD19+ proportions from all cohorts combined is a bimodal distribution that can be segregated into those consisting primarily of B cells (>50%) and T cells.

Immunophenotyping by flow cytometry of spleen cell suspensions of 345 animals demonstrated variable B and T cell proportions in all cohorts, reflecting the broad tropism of Moloney MuLV (Fig. 1d-f, gating strategy outlined in supplemental Fig. S2a). Both *BCL2* transgenic cohorts yielded a higher proportion of CD19+ B-cell lymphomas compared to wild type mice (Fig. 1g), most notably in the Vav-BCL2 cohort. Spleen suspensions segregate into two groups, with either a majority of T cells or of B cells (Fig. 1h). T-cell lymphomas were primarily CD4+, less frequently CD4-CD8-(typical of early T-cell precursor ALL), and only rarely CD8+ or CD4+CD8+ (Fig. S2b). B-cell lymphomas were generally immunoglobulin light chain positive indicating a mature B cell phenotype (Fig. S2c). MuLV infected Vav-BCL2 transgenic mice displayed a disproportionate outgrowth of PNA+ CD95+ germinal center B cells, and isotype switching to IgG as has been previously described in this strain (Egle et al. 2004)(Fig.S2c-d).

To track selection of premalignant mutations we also generated cohorts sacrificed at days 9, 14, 28, 56, 84 and 128 post-infection and harvested the spleens from these animals (Table S1). QPCR of virus transcript and copy number indicated that virus replication was detectable at day 9 and reached saturation at day 14 (Fig. 2a & b). This suggests that a high proportion of mutagenesis occurs in the first 14 days post infection, with subsequent rare events of superinfection and selective pressure shaping the mutation profile and the eventual clonal outgrowth of late-stage lymphomas.

**Figure 2.**
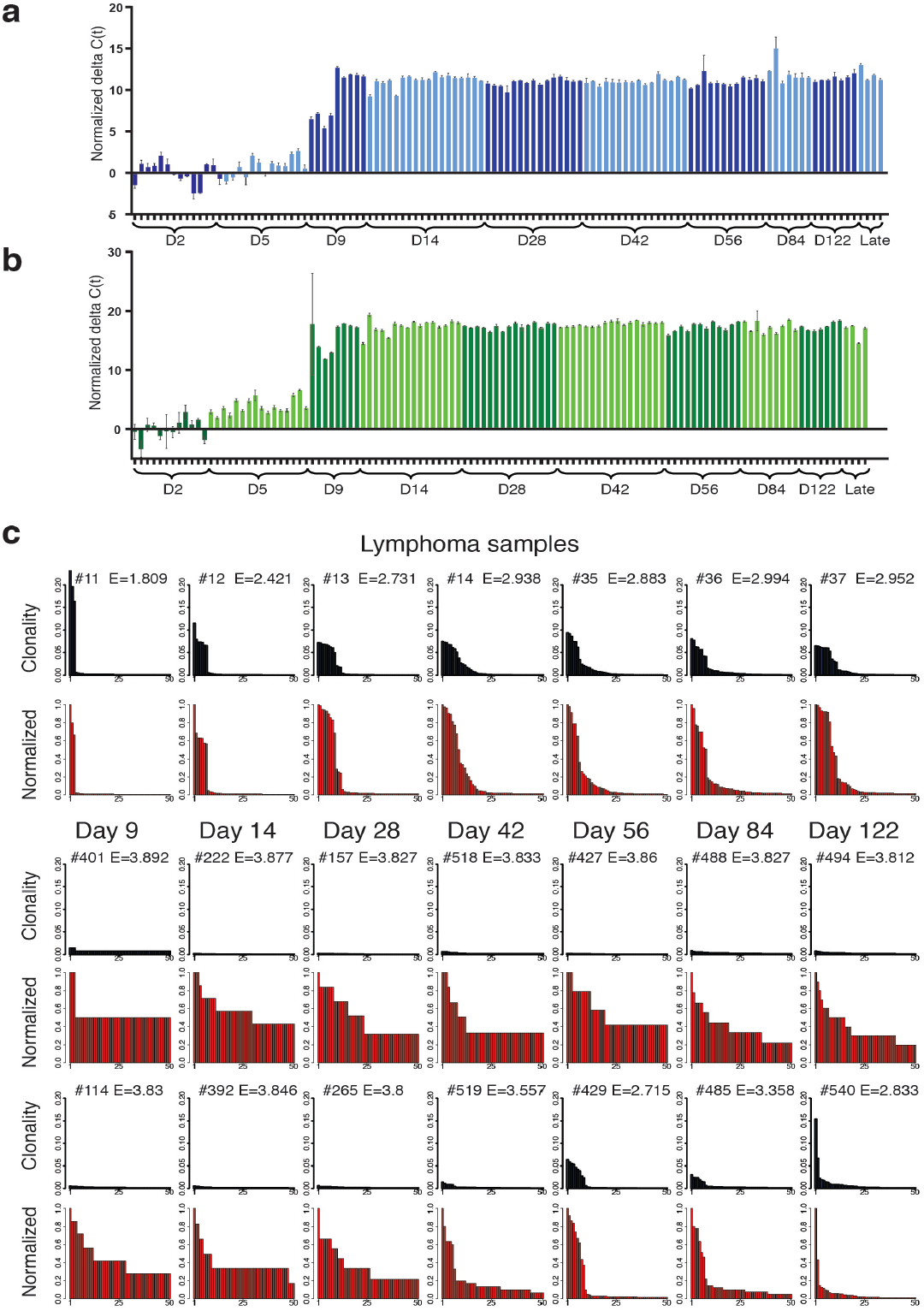
Quantifying the progression of MuLV replication and clonal outgrowth of resulting lymphoma. Virus copy number and expression level was quantified by QPCR of genomic DNA (a) and RTQPCR of cDNA (b) extracted from spleen samples of time course animals. (c) Profiles of the relative abundance of the top 50 most clonal integrations from a cross section of mature lymphoma and time course samples are represented as bar graphs. Non-adjusted clonality is indicated in blue, normalized clonality (such that the most abundant integration has a value of 1) are the graphs in red. Premalignant animals from early time points display a relatively flat profile whereas later time points and mice with symptomatic lymphoma show clear signs of clonal outgrowth. Shannon entropy values (E) are displayed on each graph.

Retroviral integration sites from all animals were identified using a novel Illumina HiSeq based protocol (a revision of methods described in Koudijs et al. 2011 and Uren et al. 2009) and summarized in Fig. S3). A series of test DNAs were used to generate multiple replicate libraries that were sequenced to saturating coverage. Measures of highly clonal mutations are reproducible when the same DNA sample is processed twice (Fig. S4a), and clonal integrations can be reproducibly detected when diluted 100 fold into a second DNA sample (Fig. S4b). Barcoded libraries were generated from the lymphoid organs of 355 diseased animals, 166 animals sacrificed at predetermined time points, and control DNAs (human and uninfected mice). Sequencing these libraries identified more than 700,000 unique integration sites, the vast majority of which are subclonal and represented by a single read. The relative clonality of integrations within each ligation was estimated using the number of individual sheared DNA fragments identified for each insert

### Quantifying lymphoma progression by clonal outgrowth

We sought to use mutation abundance to distinguish premalignant samples from rapidly dividing diseased samples. MuLV generates tumors with 100% penetrance, resulting from independent competing clones. A single clonal outgrowth of pure tumor cells containing few mutations will yield high coverage of each mutation, whereas a clonal outgrowth with dozens of concurrent mutations alongside a large proportion of non-tumor DNA will yield low coverage for even the most clonal mutation. There may also be multiple independent clones or related subclones within each animal.

As such, for comparison between samples we generated normalized clonality values (NC values) where the most clonal integration within each sample was normalized to a value of 1. In Fig. 2c the 50 insertions with the highest normalized clonality values within each sample are ordered by their relative abundance and plotted as bar graphs. Premalignant samples from the time course had a flat profile of primarily subclonal mutations with the majority represented by a single read/DNA fragment, whereas later stage lymphoma samples are dominated by highly clonal outgrowth, with between 1 and 20 clonal integrations.

We used two approaches to compare levels of clonal outgrowth. Entropy was employed as a description of the clonality of integrations within each sample (a method based on the prior use of the Shannon entropy to estimate clonal outgrowth of T lymphoma (Brown et al. 2016) and in mathematical models of leukaemia (Baldow et al. 2016)). Entropy calculations from the 50 most clonal integrations yielded high scores for premalignant samples and low scores for advanced-stage lymphoma.

As an independent approach, we used distance measures to cluster clonality profile by their shape. Dynamic Time Warping (Giorgino 2009) and the Kolmorogov-Smirnov test were both used to measure difference in shape between all insert profiles and a distance matrix was constructed from these. Clustering on these matrices yields two clusters (Fig. 3a) that place the majority of premalignant time course samples within the cluster with higher entropy values > 3.5 (Fig. 3b & c). Throughout the time course, entropy values remain high until day 56 and 84 when an increasing fraction of samples display lower values indicating clonal outgrowth (Fig. 3d). Overall, clonal outgrowth is a function of mouse age and strongly correlates with symptomatic disease. Diseased mice (i.e. those sacrificed due to symptoms) with entropy > 3.5 likely represent animals where lymphoma arose in other organs (bone marrow/thymus and lymph nodes) and had not disseminated to the spleen tissue analyzed. For subsequent analyses, we define late-stage samples as those with clonal outgrowth with a low entropy (< 3.5) and use a cut-off of entropy < 3.5 and normalized clonality > 0.1 to define late-stage clonal insertions (Fig. 3e & 3f).

**Figure 3.**
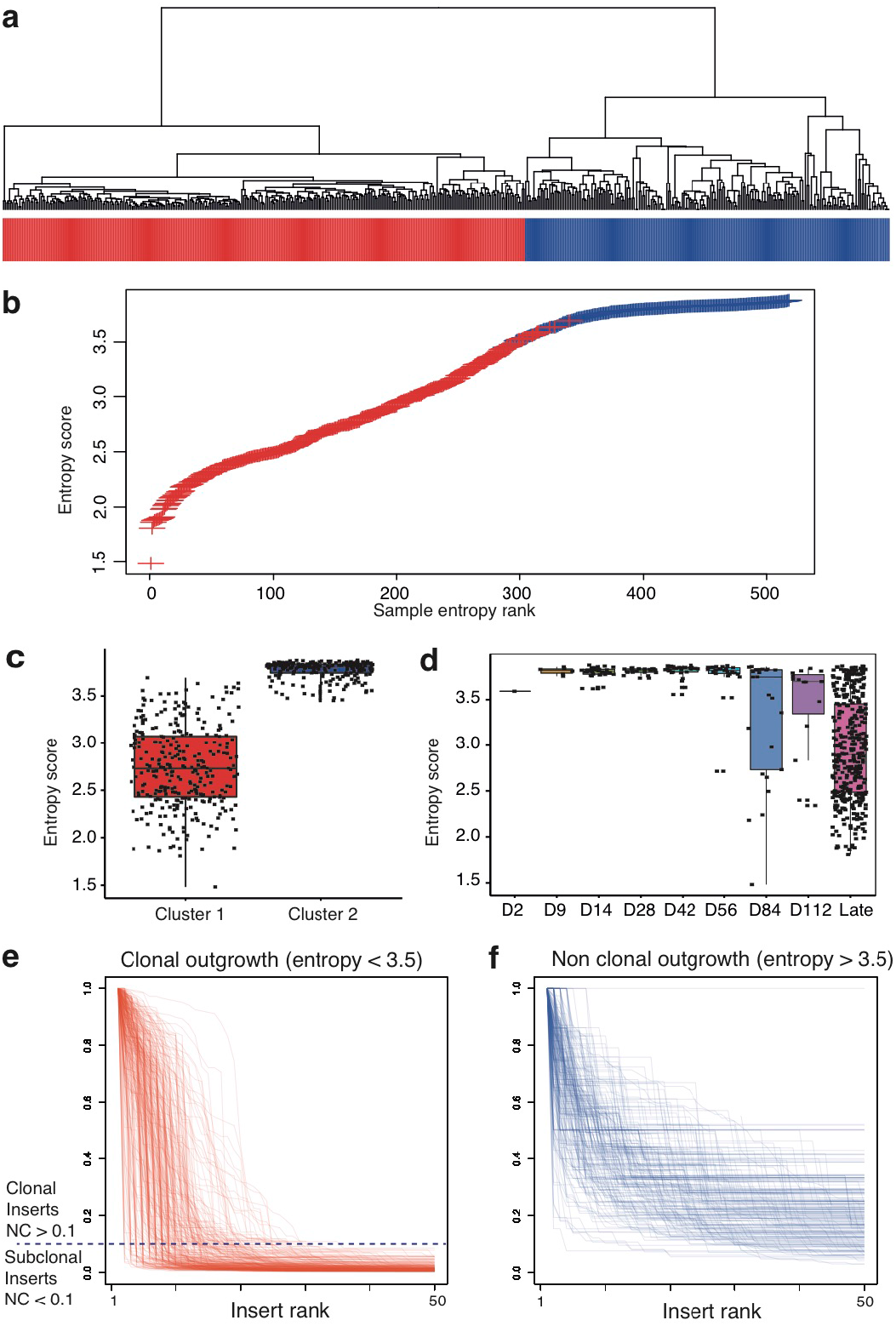
Using entropy as a measure to segregate premalignant samples from clonal outgrowth of lymphoma. a) Dynamic Time Warping was used to cluster clonality profiles of all samples and identifies two major groups; premalignant samples (blue) and samples undergoing clonal outgrowth (red). Near identical clusters were obtained using the Kolgomorov Smirnov statistic (not shown). b) Samples are plotted comparing entropy score by rank and individual samples are colored by cluster branch, indicating both entropy scores and clustering give a similar bifurcation of samples. c) Distribution of entropy scores between the two clusters indicates an entropy value of 3.5 effectively separates the groups. d) Distribution of entropy scores between different time points indicates a progressive increase in the frequency of clonal outgrowth. Superimposing the clonality profiles of all samples within each cluster indicates consistent shape within the low entropy group (e) and within the high entropy group (f). A normalized clonality value of 0.1 is used to differentiate clonal and subclonal mutations within the late stage clonal outgrowth samples.

### Kinetics of mutation selection throughout a time course

MuLV has integration biases that vary substantially throughout the genome such that mutation density is insufficient to differentiate selected driver mutations from passengers. For this reason we assessed selection over the time course to define driver mutations. First, by limiting analysis to the 3051 clonal integrations of the late-stage lymphomas we identified 311 common integration sites by Gaussian kernel convolution (de Ridder et al. 2006) (GKC) i.e. by estimating the smoothed density distribution of integrations over the genome compared to random distributions (supplemental table S2). Candidate genes were automatically assigned using the KCRBM R package (de Jong et al. 2011). Examining all insertions within 100kb windows of clonal common integration site (CIS) peaks over all time points, demonstrated a gradual increase in the proportion of inserts at these loci from day 9 through to late-stage lymphoma samples (Fig. S4a-c).

To quantify the significance of this selection we used contingency table tests (Fisher’s exact) to compare the number of integrations in windows surrounding loci in early-stage mutations (days 9 and 14), late-stage clonal, and late-stage subclonal mutations. Ranking loci using the exact test comparisons between early and late-stage mutations yields similar results using either subclonal or clonal mutations (Fig. S4d, supplemental table S2).

P-values for all early/late, clonal/subclonal comparisons for the top 50 clonal CIS loci are illustrated in Fig. 5 in the blue heat map. In some cases high ranking clonal CIS loci demonstrate weak selection between early mutations and late-stage clonal mutations (e.g. *Bzrap, Rreb1*), suggesting these are more likely to be passenger mutations resulting from integration site biases of MuLV. Late-stage subclonal mutations outnumber clonal mutations by 100 fold. Including these subclonal mutations in the analyses i.e. comparing all late-stage integrations to early mutations, reveals some CIS loci with selection that is more significant than in clonal analysis alone (supplemental table S2), including some corresponding to verified human cancer genes such as *REL, EBF1, ERG, ELF4, MYCL, KIT* and *KDR*. The finding of known cancer drivers at these loci demonstrates that this analysis of selection over a time course offers an enlarged dataset from which to identify previously validated cancer drivers and by extension, potentially identify novel genes not previously implicated in disease.

In addition to selection between early and late-stage samples, we also considered other criteria as evidence for selection. A recent study of MuLV induced T-cell lymphomas used orientation of integrations as evidence that there is selection for deregulation of nearby genes (Huser et al. 2014). The red heat map of Fig. 4 indicates that the majority of the top 50 clonal CIS loci also have a significant bias for subclonal integrations on one strand or the other. We additionally explored whether integrations also demonstrate phenotypic bias (selection specific to B-cell or T-cell lymphoma, the yellow heat map) or genotypic bias (selection in cooperation with the *BCL2* transgenes, the green heat map). The majority of the top 50 clonal CIS loci demonstrate biases of strand specificity, lymphoma subtype and/or genotype specificity. Importantly there is substantial overlap between all four selection criteria (stage, orientation, immunophenotype and genotype) suggesting these criteria can be used in concert to provide corroborating evidence for selection.

**Figure 4.**
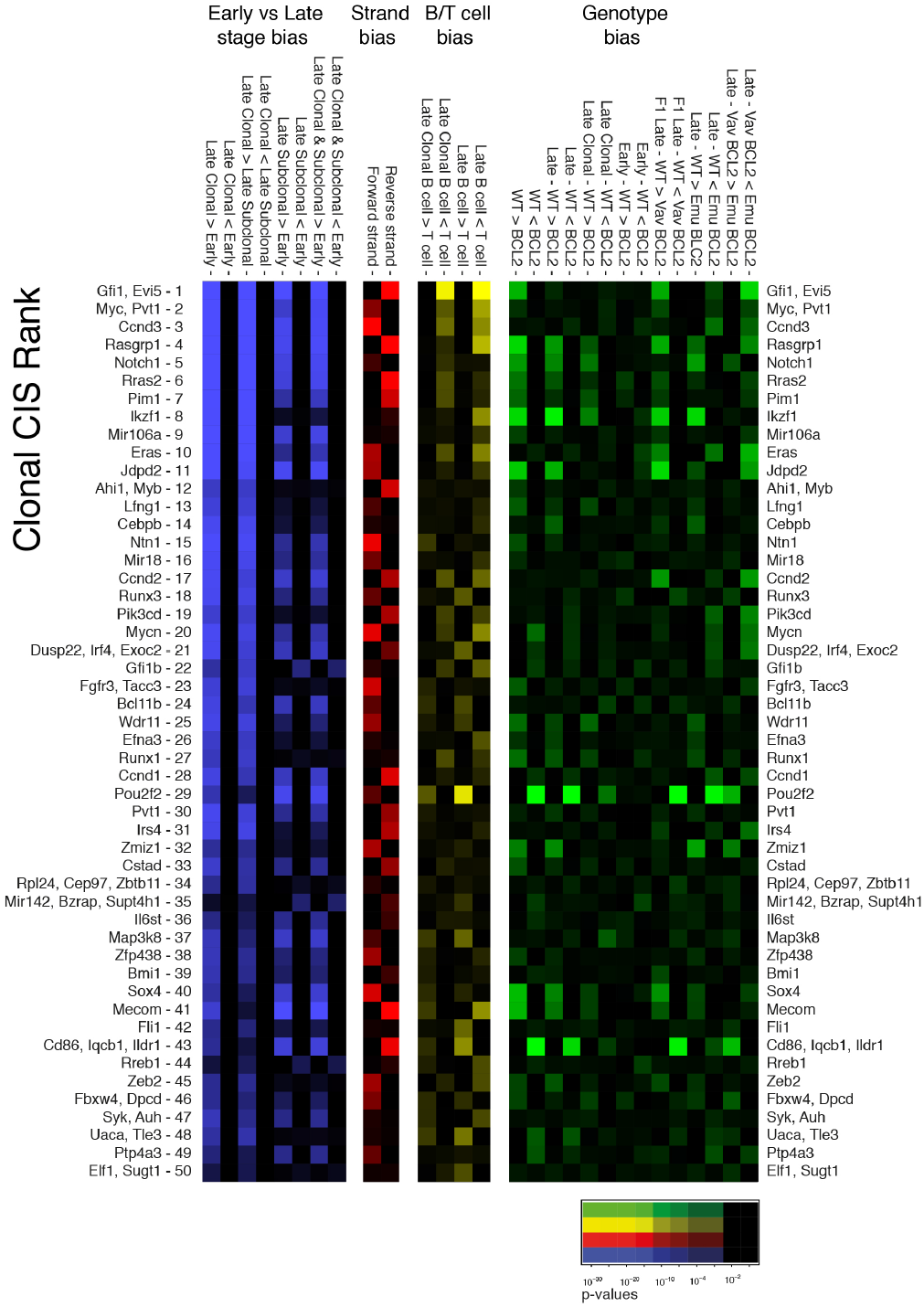
Multiple criteria indicate selection of both clonal and subclonal mutations at CIS loci. Four heat maps representing the relative levels of selection observed between different categories of integrations. Fisher’s Exact tests were performed counting the inserts within 100kb windows surrounding each of the top 50 clonal insert loci. Blue indicates comparisons between early and late stage integrations. Red represents integration orientation bias (forward or reverse strand). Yellow represents specificity for B cell (>50% CD19) versus T cell lymphomas. Green represents specificity between different genotypes. P-values for Fisher’s exact tests are indicated by color intensity.

**Figure 5.**
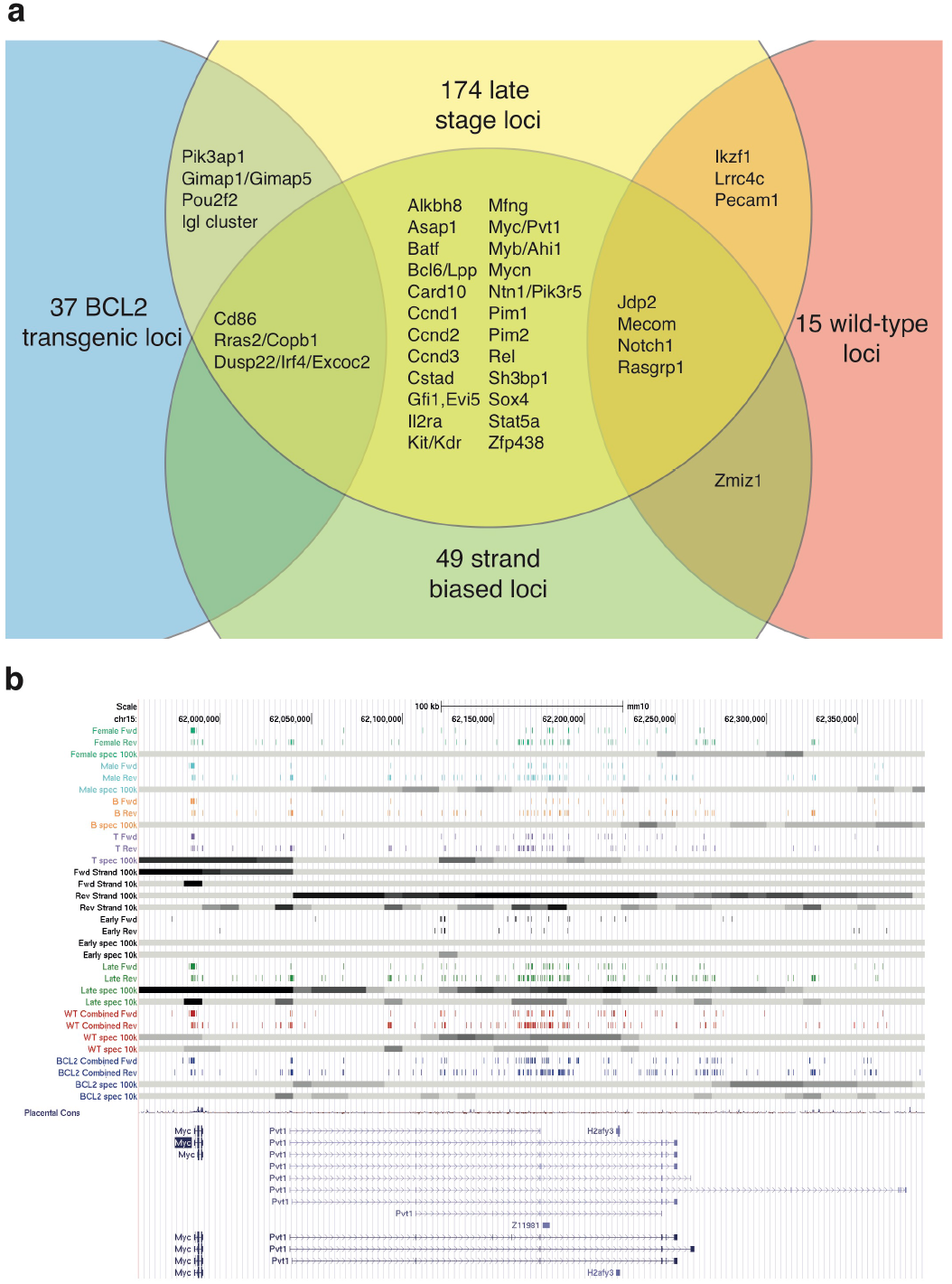
Genome wide scanning of subclonal mutation distributions identifies regions undergoing selection. a) Genome wide contingency table tests of all mutations identifies loci that are late stage specific, strand biased and genotype biased. The Venn diagram demonstrates substantial overlap between loci identified by these criteria. b) Distribution of integrations over the *Myc/Pvt1* locus. Each row of colored vertical lines represents the forward and reverse strand integrations of each category of mice. Grey bands below each colored row represent the level of selection evidenced by contingency table tests. Late stage specific integrations are evident throughout the region however integrations upstream of *Myc* are primarily on the forward strand and T cell specific whereas integrations within the *Pvt1* gene are in the reverse orientation and somewhat biased toward wild type mice.

**Figure 6.**
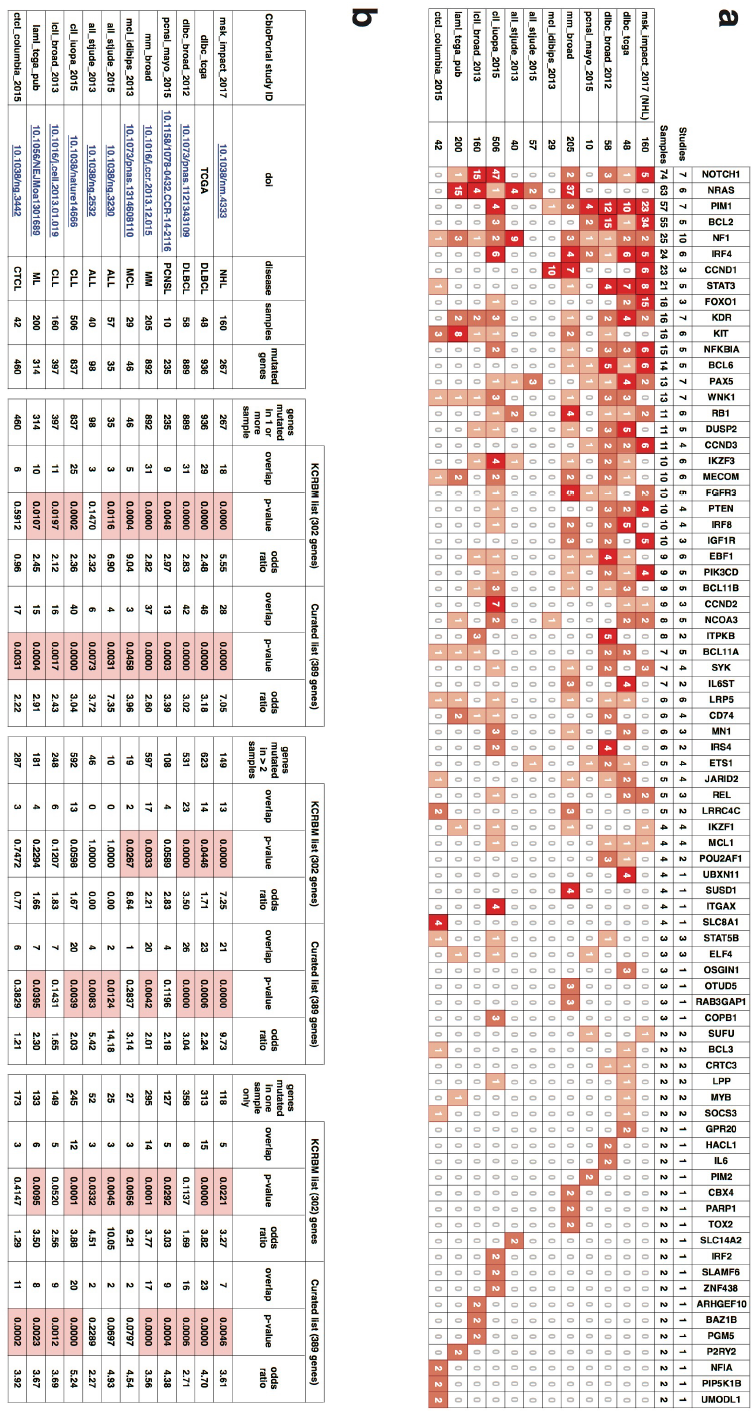
Overlap of CIS loci with exome sequencing studies of hematologic malignancies. Human orthologues were identified for all candidate genes (an automated list KCRBM, and a curated list) using biomart and compared to lists of genes with coding mutations in 12 cohorts of hematologic malignancy in cBio portal. a) All genes found mutated in at least two samples over all cohorts are listed with mutation counts from each cohort. The full overlap for all cohorts is listed in supplemental table S7. b) The significance of overlap between the set of candidate orthologues and the set of mutated genes in each study is calculated using a Fisher’s exact test.

### Identifying loci undergoing selection throughout the entire genome

The absence of clonal mutations within a region does not rule out selection in that region. Of the 300,000 integrations from late-stage diseased animals, only a fraction are located within the regions surrounding clonal CIS loci. To see if selection over the time course can identify known cancer gene loci in regions outside clonal mutation CIS loci, we extended analysis across the entire genome using both 10kb tiling and 100kb sliding windows at 10kb intervals. The distribution of integrations in different subsets of samples, across a single chromosome (chr15) is illustrated in supplemental Fig. S7a. Tracks representing inserts and the relative levels of selection across the genome (calculated using exact tests) indicates extensive selection for mutations occurring outside regions identified by GKC clonal inserts.

Examining the entire genome reveals significant local biases for early versus late-stage, strand bias, genotype and immunophenotype (B cell/T cell lymphoma) (table S3). After multiple testing correction, we identified 170 late-stage specific loci with a false discovery rate (f.d.r.) below 0.05, including dozens of windows containing only subclonal insertions. The Venn diagram in Fig. 5a illustrates that there is substantial overlap of late-stage selected loci with equivalent loci found to be strand specific (49 loci), and genotype specific (37 *BCL2* loci, 15 wild type loci). This is also true for immunophenotype specific loci (19 B cell loci, 11 T cell loci). Although biases were observed for some loci between males and females, none were found to be significant after multiple testing correction (data not shown).

In regions of high insert density subclonal mutations form a high-resolution map of the selective pressures surrounding known oncogenes. The central region of chromosome 15 (chr15:62,000,000-63,000,000) with the highest concentration of late-stage integrations is the *Myc/Pvt1* locus (Fig. 5b). The surrounding region harbors multiple clusters of selection spanning from upstream of *Myc*, and extending through multiple clusters downstream as far as the *Gsdmc* gene family locus (Fig. S7b). This distribution of selected mutations over a 2Mb region concurs with the recent finding that copy number gains of the entire segment, incorporating *Myc, Pvt1* and the *Gsdmc* family locus, is required to give acceleration of cancer in mouse models (Tseng et al. 2014).

Orientation bias is a unique criterion, in that it is independent of the integration biases of MuLV that may be influenced by cell type or genotype. To validate that orientation bias is indeed a function of selection we calculated bias using equal numbers of integrations from early and late-stage cohorts (i.e. 80,000 integrations). No loci from the early-stage inserts are significant after multiple testing correction, but the late-stage insert subset identifies 16 loci. This illustrates that the increased significance of strand bias in the late-stage cohort is not merely a function of greater statistical power from larger number of integrations, but rather evidence of selection for integrations that deregulate or disrupt genes (table S4).

### Selection of mutations effectively identifies known cancer drivers

Supplemental table S5 lists all candidate genes associated with one or more of our selection criteria. Table 1 lists the subset of these loci corresponding to genes from the cancer gene census (Futreal et al. 2004); in total 47 genes located at 43 loci. Of the 47 genes, 27 map within 200,000kb of a clonal CIS with a p-value < 0.05, however an additional 21 genes at 20 loci are implicated by subclonal selection criteria that were not identified by clonal CIS demonstrating subclonal mutations can provide additional statistical evidence to implicate cancer drivers for loci lacking sufficient clonal mutations to make this determination.

**Table 1.**
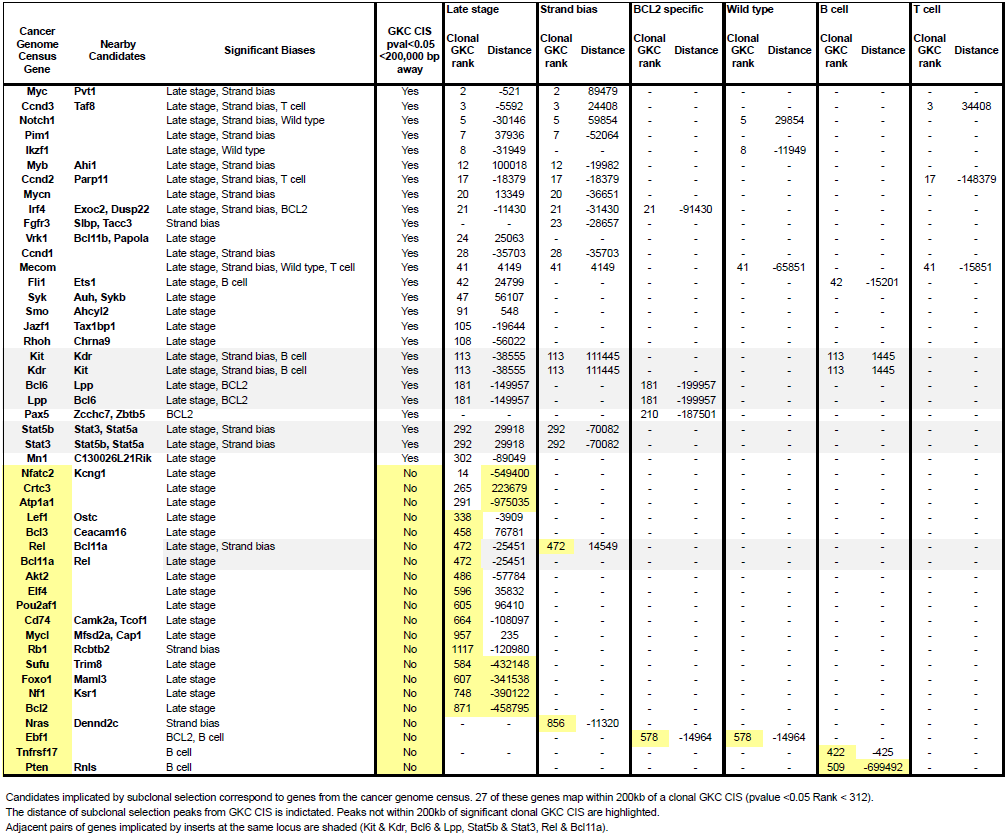
**Analysis of subclonal integrations increases the proportion of cancer gene census loci identified.** Cancer gene census genes were identified by GKC CIS analysis using late stage clonal mutations and by genome wide scanning for selection of subclonal mutations. Importantly the subclonal mutation analysis identifies known cancer drivers not identified as significant CIS by GKC of clonal mutations. Candidates implicated by subclonal selection correspond to genes from the cancer genome census. 27 of these genes map within 200kb of a clonal GKC CIS (p-value <0.05 Rank < 312). The distance of subclonal selection peaks from GKC CIS is indicated. Peaks not within 200kb of significant clonal GKC CIS are highlighted. Adjacent pairs of genes implicated by inserts at the same locus are shaded (Kit & Kdr, Bcl6 & Lpp, Stat5b & Stat3, Rel & Bcl11a).

We also compared the list of candidate genes identified by any criteria with a set of 12 cohorts of hematological malignancies present in the cBio portal (Cerami et al. 2012). All protein coding candidates generated by KCRBM (without curation) or the curated candidate gene lists were used to identify human orthologues using BioMart (http://www.ensembl.org/biomart/martview/). The set of 78 MuLV candidate genes found mutated in 2 or more samples from any study is depicted in Fig 5a. For the majority of cohorts we find significant overlap between either the KCRBM candidates or the full set of mutated genes (Fig 5b). When limiting analysis to genes mutated twice in each study we see most overlap with a pan NHL study (consisting of DLBCL and FL) and cohorts of mature B cell derived lymphoma (DLBCL, MM, MCL). Importantly the overlap is also significant when examining the set of genes mutated only once in each study i.e. the set of most rarely mutated genes from most cohorts overlaps significantly with the candidate lists, demonstrating this dataset can be used as corroborating evidence for rarely mutated genes in human sequencing cohorts.

Across the set of genes identified there is a notable prevalence for genes that are known to be deregulated by translocations and/or copy number aberrations as well as those found in genome wide association studies in humans. Using a list of 278 selected regions identified in this screen by any criteria, 273 were mapped unambiguously to orthologous region on Hg19. We overlapped this set of loci with focal copy number aberrations of 5 human studies of mature B cell lymphoma and found a significant degree of enrichment in 4 of the 5 datasets (table S8).

Recurrent large-scale copy number changes in human B NHL suggest the involvement of multiple genes within these regions. We find a number of corresponding loci where selection is evident over large regions incorporating multiple genes. Aside from the abovementioned *Myc/Pvt1/Gsdmc* family locus (Fig. S7b) we see multiple selected regions surrounding the *Rel/Bcl11a* locus (orthologous to human 2p12-16 amplicons of CLL & DLBCL Fig. S7c), the *Slamf* gene family which regulate lymphocyte survival, activation and co-stimulation (orthologous to human 1q21-23 amplicons of multiple myeloma and DLBCL Fig. S7d) and the *Gimap* gene family of GTPases that also regulate lymphocyte survival and development (orthologous to amplicons of the distal arm of human 7q seen in FL, DLBCL and Burkitt lymphoma) (Fig. S7e). We also see multiple selected loci spanning the region surrounding *Prdm1* (orthologous to deletions of human 6q21 in B-cell NHL and other hematologic malignancies Fig. S7f).

The use of *BCL2* transgenic animals expands the scope of mutations identified beyond loci typically identified by MuLV in wild type animals. Of 37 loci that are *BCL2* transgene specific, and 19 loci that are B cell specific, we find selected regions near known B-cell lymphoma and leukemia drivers including *Pou2f2, Ebf1, Ikzf3* and *Bcl6*. The most specific locus for both *BCL2* transgenic animals and for B cell lymphoma is *Pou2f2* (Fig. S7g) which is recurrently mutated in human FL and DLBCL. Recurrent missense mutations reduce Pou2f2 transactivation activity and B lymphoma cell lines expressing these have a survival advantage (Li et al. 2014), conversely DLBCL cell lines appear to be addicted to *POU2F2* expression (Hodson et al. 2016). The position of insertions within several known tumor suppressor genes is suggestive of disruption. Similar intragenic distributions of integrations have previously been described for the tumor suppressor *Ikzf1* which is mutated or deleted in both B and T-ALL (Gupta et al. 2016; Mullighan et al. 2008) (Fig. S7h), and this pattern is also observed for the tumor suppressors *Ikzf3* and *Ebf1* (Fig. S7i & S7j), both of which have inactivating mutations in FL and DLBCL but more typically B-ALL (Morin et al. 2013; Gupta et al. 2016; Bouska et al. 2014; Mullighan et al. 2007). *Cxxc5* (late-stage specific) also has a similar pattern of intragenic integrations (Fig. S7k) and is deleted and epigenetically silenced in acute myeloid leukemia (Kühnl et al. 2015). The majority of our insertions at the *Pou2f2* locus are intragenic, consistent with the tumor suppressor function observed in FL.

To find additional evidence supporting the role of candidate genes in hematologic malignancies, we conducted an extensive review of the literature, with an emphasis on data from human lymphoid malignancies and a particular focus on *BCL2* driven B cell lymphoma. Supplemental table S6 lists evidence for 194 genes from our list of candidates flanking selected loci. In addition to 47 genes identified in the cancer genome census we also find 30 genes identified in the Intogen mutational cancer driver gene list (Gonzalez-Perez et al. 2013).

### Selection of loci implicated by GWAS of lymphoid malignancies

In a previous study we found overlap between CIS loci and the set of loci associated with familial chronic lymphocytic leukemia (CLL) (Kool et al. 2010). Supplemental table S9 summarizes the recent literature of GWAS studies of ALL, FL and DLBCL and identifies overlap with the set of candidate genes. The *IKZF1* and *PIP4K2A*/*BMI1* loci are associated with ALL (Xu et al. 2013; Moriyama et al. 2015). GWAS of mature B cell lymphomas have identified associations with follicular lymphoma (*LPP, HLA* loci, *PVT1, CXCR5, ETS1* and *BCL2*) and with DLBCL (*LPP, EXOC2, HLA-B and PVT1*) (Cerhan et al. 2014; Bassig et al. 2015).

The second most specific locus for *BCL2* transgenic mice is *Cd86* (late-stage, strand bias, BCL2) which is suggestively implicated by two GWAS studies of FL and DLBCL (Skibola et al. 2014; Cerhan et al. 2014; Bassig et al. 2015). We additionally find other loci encoding co-stimulatory/co-inhibitory signaling (Fig. S7l-p). The loci encoding *Cd86* ligands *Ctla4* and *Cd28* and their neighboring homologue *Icos* show late-stage specific selection, and polymorphisms in this region have also been associated with various NHL subtypes (Piras et al. 2005). Other members of the B7 family and their receptors are also implicated by insertions including *Icoslg* (late-stage) and the *Cd274* (Pdcd1lg1) and *Pdcd1lg2* locus (late-stage with f.d.r. = 0.081) as is their receptor *Pdcd1* (late-stage). *Pdcd1lg2* is amplified and rearranged frequently in primary mediastinal large B cell lymphoma (Twa et al. 2014) and increased expression is speculated to inhibit anti lymphoma T cell responses (Shi et al. 2014).

Late-stage biased insertions are located near the *H2-D/H2-Q* locus (orthologous to the MHC I *HLA-B/C* loci) and *BCL2* transgenic biased clusters are found near the MHC Class II beta chains *H2-Ob, H2-Ab1, H2-Eb1* and alpha chains *H2-Aa, H2-Ea-ps* (orthologous to the MHC II *HLA-DRB/HLA-DQB loci*) (FigS7q). Aside from the abovementioned GWAS associations both regions are deleted in human DLBCL (Booman et al. 2008; Monti et al. 2012; Sebastián et al. 2016). There are also suggestive clusters of late-stage insertions surrounding the MHC Class I components *H2-T24/T23/T9/T22/BI/T10/T3/Gm7030* (equivalent to the HLA-E locus) and the MHCII alpha/beta chains *H2-Oa/H2-DMa/H2-DMb* (equivalent to the *HLA-DOA/HLA-DMA/HLA-DMB* region), suggesting roles for both classical and non-classical MHC components in lymphoma progression.

### Negative selection of mutations throughout lymphoma development

Intriguingly we observed many loci throughout the genome that are early-stage specific i.e. undergoing negative selection between early and late-stage cohorts, suggesting these integrations become detrimental to survival and expansion of developing lymphoma cells. The most significant of these is the *Smyd3* locus where a cluster of intragenic insertions surrounding exons 6-8 are present to a significantly lesser extent in the late-stage lymphomas samples (supplemental figure 7r). SMYD3 is a methyltransferase that methylates H3K4 and H4K5 and over-expression has been observed in a variety of tumor types. SMYD3 methylation of MAP3K2 activates MAP kinase signaling and loss of *Smyd3* delays development of both pancreatic and lung tumors (Mazur et al. 2014). Presumably integrations that disrupt *Smyd3* expression are detrimental and hence selected against in lymphomagenesis. *SMYD3* loss potentiates the effects of MEK1/2 inhibition on tumor growth (Mazur et al. 2014) and SMYD3 inhibitors have been shown to inhibit the growth of tumor cell lines (Peserico et al. 2015). The example of the *Smyd3* locus demonstrates the potential for time course mutation analysis to not only identify cancer drivers, but also potential targets that whilst not mutated, are essential for tumor cell growth.

### Co-mutation analyses using subclonal mutations

Understanding which genes cooperate in lymphomagenesis can inform the biology and subtype of disease, however co-mutation analyses of subclonal mutations is complicated by the potential presence of multiple independent subclones. To minimize these effects, we performed contingency table tests to identify co-mutation rates, limiting analysis to the most clonal integrations (NC>0.1), increasing the likelihood that mutations are present in the same clone. Rather than discard all subclonal insertions, we additionally performed analyses looking for associations of clonal mutations with the subclonal mutations, based on the assumption that subclonal mutations may be present in the same cells as clonal mutations from the same sample. Both approaches yield overlapping results overall, but the latter greatly improves the statistical power of the associations identified (supplemental Fig.S8).

We have previously demonstrated that co-mutation analysis can be compromised by the pooled analysis of phenotypically and genotypically distinct groups, which creates false positives from genes that are co-mutated or mutually exclusive due to a primary association with tumor subtype rather than other mutations (Kool et al. 2010). For this reason we devised an online tool that allows subsets of tumors with restricted phenotypes, genotypes and mutation profiles to be queried http://mulv.lms.mrc.ac.uk/coocc/index.php.

## DISCUSSION

In this study we present the most comprehensive analysis of MuLV-driven lymphomagenesis produced to date, identifying 700,000 mutations from 521 infected animals with an average of more than 1000 subclonal mutations per sample. By developing a framework that incorporates subclonal mutation frequencies in both premalignant cells and late-stage tumors, we enhance the statistical power enabling the identification of known driving events in cancer and further implicate dozens of novel loci in the biology of lymphomagenesis, in some cases independently of evidence of clonal expansion.

The resolution of mutation coverage illustrates considerable complexity in the position and orientation of selected integrations in the vicinity of verified cancer drivers, and suggests uncharacterized locus-specific mechanisms by which these mutations modify expression in a position dependent manner. The online repository (http://mulv.lms.mrc.ac.uk) allows researchers studying lymphoid malignancies to query custom subsets of data for genome wide associations of a gene of interest with tumor type and mutation status, and create custom tracks for the UCSC genome browser (Kent et al. 2002). Tracks on the UCSC genome browser can also be browsed to examine the selection biases at specific loci of interest http://mulv.lms.mrc.ac.uk/ucsc/index.php) and subsets of tumors can be queried to identify co-mutated genes within phenotypically/genotypically matched lymphomas.

Recent whole genome sequencing of cohorts of hundreds of patient samples illustrates the challenges of identifying driver mutations outside the exome (Puente et al. 2015; Nik-Zainal et al. 2016). Recurrent clonal mutations follow a power law distribution, with statistically intractable rare events making up the bulk of mutations in many tumor types. Proving which of these contribute to disease remains a bottleneck that can only be partly alleviated by larger cohort sizes. Whilst the clonal mutations in MuLV driven tumors match a similar power law distribution, selection of mutations identified at the subclonal level strongly correlate with clonal mutations and/or known cancer drivers, suggesting that these mutations can provide statistical support for the role of rarely recurrent clonal mutations. Reanalysis of existing cohorts of tumors from other insertional mutagenesis screens alongside equivalent premalignant tissue may greatly expand the yield of cancer drivers identified as well as eliminate false positives.

Many tumor types display background mutation rates that vary throughout the genome. For instance, mature B cell lymphomas are in part driven by aberrant somatic hypermutation (Khodabakhshi et al. 2012). This variation can confound the identification of driver mutations outside non-coding regions. The overlap we find between independent criteria as evidence of selection (disease stage, tumor type, genetic interactions and strand bias) using a mutagen that exhibits strong regional variation in distribution, demonstrates it is possible to use these criteria as a mitigant of regional variation in mutation frequencies. Furthermore, visualizing this selection as a continuum at multiple scales using multiple parameters allows intuitive differentiation of recurrent selection in non-exonic regions from mutation hotspots.

Insertional mutagenesis screens complement the characterization of human tumor genomes identifying genes that play a crucial role in tumor biology even where they are not subject to exonic mutations, frequent translocations, or focal copy number changes. Loci identified in this and previous MuLV screens correspond to NHL cancer drivers, particularly with copy number aberrations, translocations and loci identified by GWAS. This is consistent with MuLV primarily acting via deregulated expression of open reading frames, with a subset of loci having disrupted/truncated open reading frames. Proving that deregulated but intact open reading frames are cancer drivers is problematic, particularly for tumor types such as CLL where the number of coding mutations per cancer genome is low and substantial epigenetic deregulation has been observed (Guièze and Wu 2015). Aside from the candidates we find with a supporting role in the literature, this study implicates hundreds of other candidates with equal significance that can be used as a resource to prioritize the study of human cancer drivers and potential therapeutic targets.

Targeted resequencing of recurrently mutated genes in CLL has demonstrated that coding subclonal mutations also undergo significant selection and even convergent evolution (Jethwa et al. 2013). Currently selection of subclonal mutations in cancer is difficult to prove outside the coding regions of known cancer drivers, in part because the error rates of existing NGS platforms limit detection of single nucleotide allele frequencies to > 1%. Novel technologies for detection of lower abundance mutations are in development (Chen-Harris et al. 2013; Gerstung et al. 2012; Schmitt et al. 2012), although their throughput and coverage is limited. This study demonstrates the value of applying genome wide, subclonal mutation detection to large cohorts of cancer genomes. Analyzing thousands of subclonal mutations per sample, in both malignant and premalignant tissue, and incorporating these into a framework that combines tumor genotype and phenotype, not only provides supporting evidence for rarely mutated cancer drivers, but also potentially widens the spectrum of genes encoding therapeutic targets.

## METHODS

### Animal work

All procedures were performed in accordance with the UK Home Office Animals (Scientific Procedures) Act 1986. BCL2-22 (B6.Cg-Tg(BCL2)22Wehi/J, http://jaxmice.jax.org/strain/002318.html) were bred with wild-type C57BL/6 and BALB/c mice (Charles River, UK). C57BL/6 Vav-BCL2 mice were bred with wild type BALB/c mice to produce (BALB/c x C57BL/6) F1 Vav-BCL 2 mice.

MuLV was prepared by transfection of 293T cells with the plasmid pNCA (Colicelli and Goff 1988) (provided by Stephen Goff, Addgene 17363). Newborns were injected intraperitoneally with 50μl MuLV supernatant. Mice were weighed weekly and monitored three times per week for signs of illness. Mice (infected and matched controls) in the time course cohort were sacrificed and lymphoid organs harvested at predetermined time points, prior to disease onset (9, 14, 28, 42, 56, 84 and 112 days). Survival cohort mice were sacrificed upon developing advanced symptoms of lymphoma and lymphoid organs were harvested and snap frozen in liquid nitrogen immediately. Cell suspensions of spleen tissue were prepared in all cases using the gentleMACS Dissociator (Miltenyi Biotec) set to programme m_spleen_1.01.

### Flow Cytometry

Cryopreserved spleen suspensions were defrosted and washed twice in buffer, PBS-2% FCS and incubated with 2.0μg Fc block per 10^6^ cells for 15 minutes. The samples were then incubated with the antibody cocktails for 15 minutes after which they were washed. The majority of samples were processed using the Attune NxT Acoustic Focusing Cytometer, Life Technologies, and the remainder were processed on a BD LSRII. All analyses were performed using FlowJo v10.2.

### Integration site cloning and GKC CIS identification

To clone virus integrations we integrated the method of Koudijs et al (Koudijs et al. 2011) and Uren et al (Uren et al. 2009) and modified these for the Illumina platform. A detailed protocol is provided in the supplemental methods document.

### Entropy quantitation

Clonality values among the early-stage samples are relatively uniform, whilst late-stage samples present few integrations with very high clonality values and most with low clonality values. To quantify this difference and thus be able to order samples from pre-malignancy to late-stage lymphoma we used Shannon entropy (Shannon 2001). The 50 highest clonality values *c*_1_, *c*_2_,…,*c*_50_ were transformed into probabilities *P*_i_:

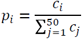

The Shannon entropy *E* over a set of probabilities *P*_1_, *P*_2_,…, *P*_*n*_ is defined as:

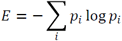

The entropy quantifies the spread of a distribution: it is zero when a single *P*_*i*_ is equal to one and all others are equal to zero; and reaches its maximum value when the probabilities are uniformly distributed (*p*_*i*_*=1/50* for every *i*). Probabilities from early-stage samples are closer to a uniform distribution and therefore the samples will have high entropy values, while the probabilities from late-stage samples are closer to a spike, providing low entropy values.

### Clustering

Samples were clustered based on the shape of clonality profiles. The top 50 ranked clonal insert NC values of each sample were compared to all other samples using dynamic time warp (dtw package) or the Kolmorogov-Smirnov test (ks.dist() function) as a measure of difference between samples (with the R package, function dist() takes as input the list of NC values and the method chosen -“DTW”-). A distance matrix that was clustered using the “average” linkage method (using the R hclust() function). Different linkage methods gave similar results (data not shown).

### Statistical analysis

#### Genome wide scanning for selected insertions

A scanning 100kb window is moved across the genome in increments of 10kb. For each window the number of insertions in each class (early/late, forward strand/reverse strand, BCL2 transgenic/wild type) is counted and the likelihood of this distribution between groups is estimated using two-tailed Fisher's exact test. By comparing neighboring windows, p-value minima are identified (i.e. windows where the p-value is higher on either side). Where minima are less than 100,000bp from each other the position with the lowest p-value is kept and others discarded. To estimate false discovery rates all insert/group assignments are randomized (e.g. the same number of inserts are early/late but the assignment is random). Local p-values are calculated and p-value minima are identified. 1000 permutations are used to determine the rate at which each p-value is identified by chance.

#### Cohort comparisons

Survival comparison of cohorts was performed using Prism 6. Significance of the differences in the proportion of B cells in cohorts was determined using Prism 6 student’s t-test.

### Data Accession

Paired mapped reads for the sequencing are available from the NCBI sequence read archive https://www.ncbi.nlm.nih.gov/Traces/study/?acc=SRP110741.

## ACKNOWLEDGEMENTS

Charles Bangham, Suzanne Turner, Jesus Gil, Vincenzo de Paulo, Niall Dillon, Matthias Merkenschlager, Amanda Fisher provided helpful comments on the data and manuscript. Andreas Villunger provided Vav-BCL2 transgenic animals. Emma Mustafa, Aleksandra Czerniak and Katie Horan assisted with animal care and monitoring. Andre Brown suggested Dynamic Time Warp as a measure of clonality profile similarity.

## AUTHOR INFORMATION

### Contributions

Conception and design: P. Webster, A. G. Uren.

Development of methodology: P. Webster, K. Takacs, B. Iadarola, J. Caceres, B.J. Bolt,

J. Kaczor, L. Game, J. Elliott, K. Naresh, A.G. Uren

Acquisition of data: P. Webster, J.C. Dawes, H. Dewchand, J. Kaczor, A.G. Uren

Analysis and interpretation of data: P. Webster, J.C. Dawes, H. Dewchand, B. Iadarola,

J. Caceres, B.J. Bolt, J. Kaczor, G. Dharmalingam, A. Paccanaro, A.G. Uren

Writing, review, and/or revision of the manuscript: P. Webster, J. Dawes, K. Takacs, B.

Iadarola, J Caceres, L. Game, J. Elliott, A. Paccanaro, A.G. Uren

Technical and informatics support (mouse maintenance, flow cytometry, reagent generation, database construction and maintenance): P. Webster, J.C. Dawes, H. Dewchand, K. Takacs, B. Iadarola, B. J. Bolt, J. J. Caceres, J. Kaczor, L. Game, T. Adejumo, J. Elliott, G. Tan, G. Dharmalingam

### Accession numbers

The sequence reads mapped to mouse build mm10 are available in the NCBI sequence read archive https://www.ncbi.nlm.nih.gov/Traces/study/?acc=SRP110741

